# BioImageIT: Open-source framework for integration of image data-management with analysis

**DOI:** 10.1101/2021.12.09.471919

**Authors:** Sylvain Prigent, Cesar Augusto Valades-Cruz, Ludovic Leconte, Leo Maury, Jean Salamero, Charles Kervrann

## Abstract

Open science and FAIR principles have become major topics in the field of bioimaging. This is due to both new data acquisition technologies that generate large datasets, and new analysis approaches that automate data mining with high accuracy. Nevertheless, data are rarely shared and rigorously annotated because it requires a lot of manual and tedious management tasks and software packaging. We present BioImageIT, an open-source framework for integrating data management according to FAIR principles with data processing.

## Main text

Advances in biological imaging over the past 20 years have been accompanied by a multitude of computerized approaches for indexation, reconstruction, processing, analysis, and exploitation of image data. As they address a variety of biological models at diverse scales of life, they are often dedicated to target case-studies (single cell scale, tissue-scale, embryos…). Consequently, imagining and developing a unified solution able to tackle any scenario in biological imaging very challenging. For more than ten years, several architectures and tools have been proposed to surmount many technological difficulties. The existing software platforms (e.g., Fiji^1^, Icy^2^, Knime^3^), developed in various programming languages (e.g., java, c++, python) so far, are unfortunately not interoperable. Additional efforts in scripting and codes development are generally required to gather some heterogeneous image processing components in ad-hoc workflows.

Meanwhile, data is hosted in storage servers associated with a database system of their own (e.g., OMERO^4^), without much possible interaction with processing and analysis tools. A more integrative approach of organizing, viewing, and analyzing data, is more and more required in data science to avoid manual management or script writing which can be tedious for non-expert scientists. Also, the management of often-massive datasets requires dedicated expertise in computer science to scale up the storage and computing resources. Finally, with the emergence artificial intelligence (AI) such as deep learning in bioimaging (e.g. DeepImageJ^5^, ImJoy^6^), the automation of processes and the implementation of processing pipelines must take all stages of the data life cycle and new human-machine interactions into account. As such, data management and processing/analysis must meet quality criteria ensuring identification, accessibility, and interoperability of the data with their processing, storage, and analysis, and finally, anticipation of future reuse. These “FAIR” principles (findable, accessible, interoperable, and reusable)^7^ impose new procedures and ethical issues to scientists whose research relies heavily on biological imaging, leading to a change in paradigm for image-based production of knowledge.

To address all these requirements, we developed BioImageIT, a unique open-source system integrating image data management and analysis, and an operational solution for handling large image data sets to the requirements of open science (Fig. 1a). In BioImageIT, data is automatically annotated and processed in a single framework, and once processed. Unlike previous solutions, BioImageIT integrates any existing data management and image processing software. For instance, data can be hosted in OMERO^4^ database using Bio-Formats^8^ and processed by applying machine learning algorithms embedded in the TensorFlow^9^ framework.

**Figure 1.**
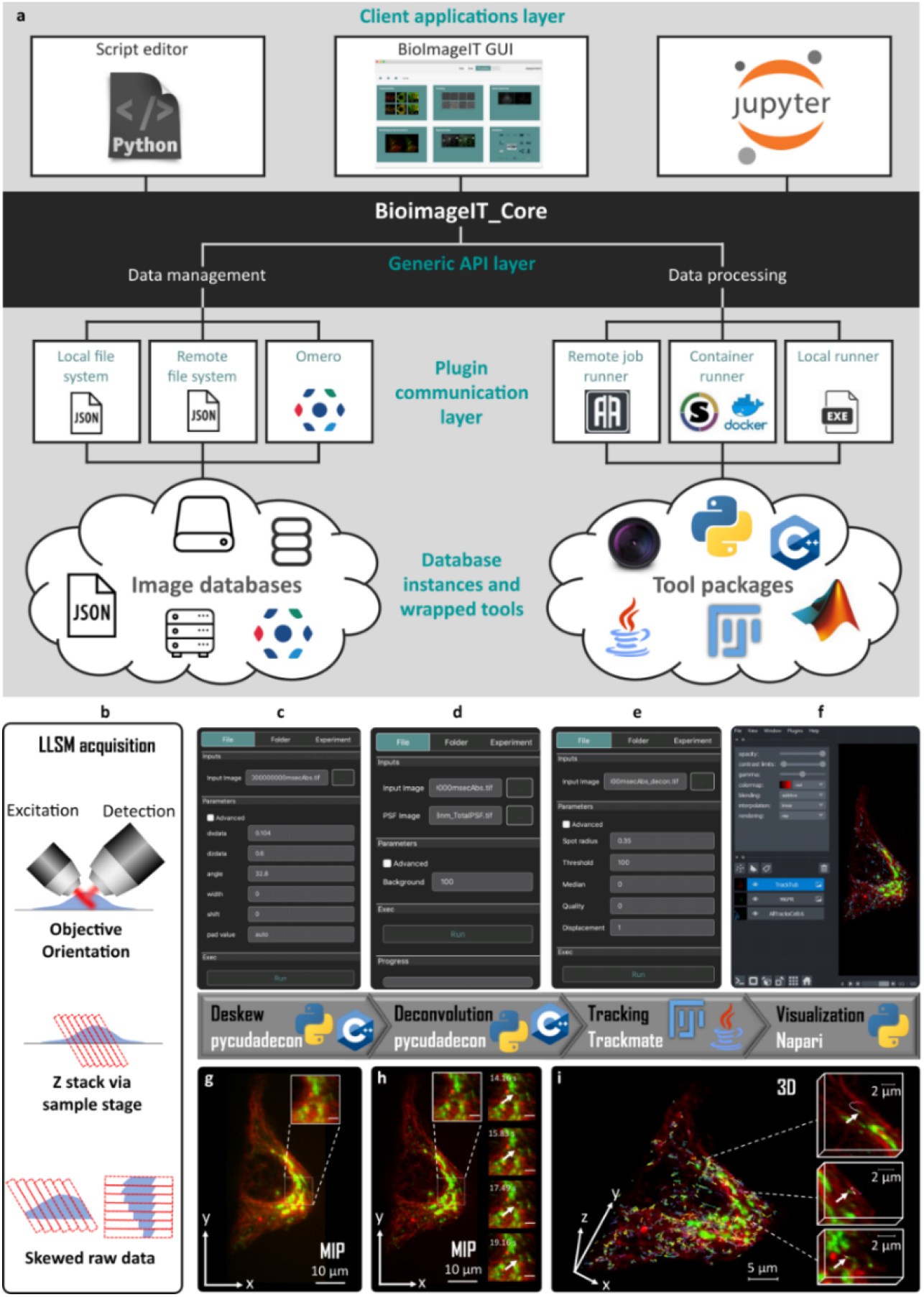
BioImageIT overview. **a**, Schematic view of BioImageIT architecture. BioImageIT core is composed of data management and data processing functionalities. Users can access plugins by a script editor, Jupyter^10^ or BioImageIT graphical interface. Data management functionalities exploit local files, remote files or OMERO database. Data processing can perform computations in remote job, container, or local runner. Image analysis is provided by plugins, potentially written in different programming languages. Developers can implement their own plugins in BioImageIT and design their Graphical Interface. **b-i**, Example of Lattice Light Sheet^11^ workflow of 3D reconstruction and tracking of live Genome-edited Hela cell (CD-M6PR-eGFP) stained with Tubulin TrackerTM Deep Red for Microtubules (see Methods). **b**, Due to the geometry of LLS scanning, 3D raw images are skewed. **c, g**, First, realignment (deskew) of raw stacks was performed using Pycudadecon (https://github.com/tlambert03/pycudadecon). **d, h**, Richardson Lucy deconvolution^12^ is performed by using Pycudadecon. **e**, CD-M6PR-eGFP vesicles are tracked by using Trackmate^13^. **f, i**, Deconvolved stacks and tracks are rendered by using Napari^14^. Full workflow, including Pycudadecon, Trackmate^13^ and Napari^14^, was gathered in BioImageIT.

Unlike previous frameworks (e.g., Galaxy Project^15^, Snakemake^16^, BIAFLOWS^17^), BioImageIT enables to address the following end-user’s issues:

### BioImageIT reconciles data management and data analysis in a common and interactive framework

Most open-source bio-image software are developed separately and specialized in either data management or data analysis. Therefore, users need to write ad-hoc scripts or to apply manual operations to process the data. BioImageIT allows importing and tagging data for each experiment. Each operation automatically generates metadata that allows to keep track of any processing or analysis and then complies with the recommended FAIR principles^7^ in data science.

### BioImageIT is interoperable and reusable

The processing tools are packaged in Docker^18^ containers or Conda^19^ packages. Users can then create data analysis workflows with software developed in any language (Supplementary Note 2). Processing tools and workflows are stored in public repositories and can be reused by another user. BioImageIT has been tested on different operating systems, including Windows 10 (Supplementary Video 1), Linux Ubuntu 21.04 (Supplementary Video 2) and MacOS Catalina 10.15.11 (Supplementary Video 3).

### BioImageIT is developer-friendly (Supplementary Note 1)

BioImageIT allows data scientists to more easily distribute new tools embedded in a Docker image or Conda package. A basic configuration file (wrapper) is only required for identification in BioImageIT

### BioImageIT is user-friendly (Supplementary Note 2)

BioImageIT consists of 3 layers (backend plugins, python API, Graphical User Interface (GUI)). Users can choose the most appropriate level of interaction with BioImageIT. Biologists may prefer to be assisted step-by-step; GUI is then appropriate for data management and analysis. Data analysts are more familiar with scripts and Jupyter^10^ notebooks using the python API. Finally, data scientists may adopt the packing backend system to provide a standalone demonstration of the new processing tool.

In summary, BioImageIT is a generic framework for data management, analysis, and traceability. Unlike previous software platforms, it meets the needs of end-users and is a very flexible solution to link annotated data and processing tools. Furthermore, it facilitates interactions between experimental and data scientists. As BioImageIT is built upon existing technologies, it may be considered as a computer overlay allowing a user-friendly use of existing software by end users, without hindering the addition of new analytical methods by more experienced developers. Finally, the ability of BioImageIT to plug in data interaction tools allows for the deployment of highly specialized instances of BioImageIT for a specific application domain, as illustrated in Figure 1 depicting Lattice Light Sheet Microscopy^11^ Data processing (Fig. 1b-i), reconstruction of Multi-Angle TIRF Azimuthal^20^ (Fig. 2), as well as advanced image processing using conventional techniques (e.g. ND-SAFIR^21^) and deep learning (e.g. N2V^22^) (Supplementary Fig. 1 et Supplementary Video 4). These examples show the ability of BioImageIT to exploit advanced imaging analysis tools and workflows applied to datasets from various advanced microscopies. BioImageIT is being deployed on several imaging platforms, members of the France BioImaging infrastructure. On this basis, our next step is to create a BioImageIT community. Documentation, source code and video tutorials are available at https://bioimageit.github.io, and help is available via GitHub https://github.com/bioimageit.

**Figure 2.**
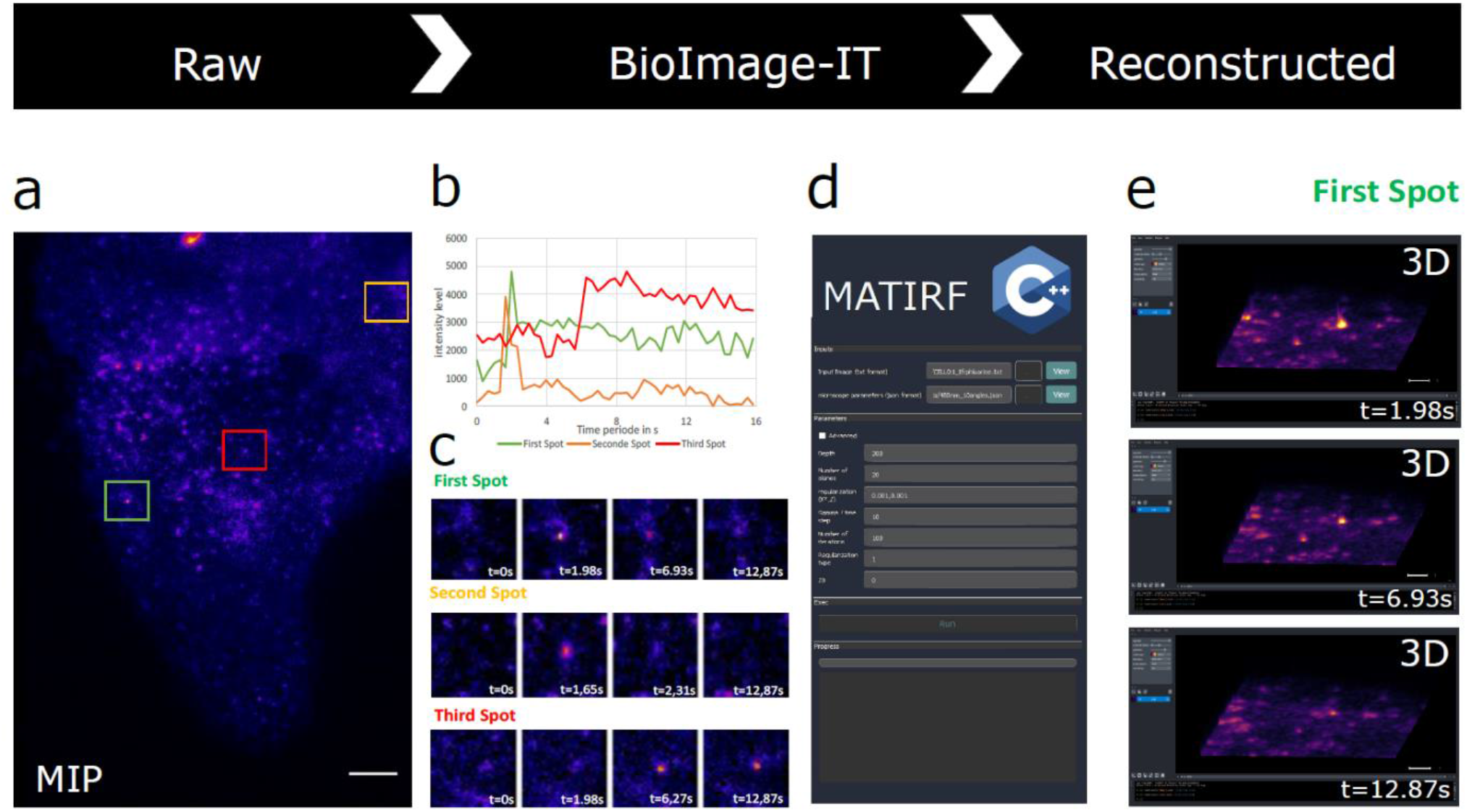
BioImageIT a method for advanced reconstruction TIRF Azimuthal Multi-Angle^20^. Live cells RPE1 cells were transfected for Transferrin Receptor-Phluorine as indicated in Methods before acquisition with Multi-Angle Ring TIRFM. **a**, Maximum intensity projection of an image stack constituted of ten angles were acquired at 30ms exposure time per angle. Scale Bar 5µm. **b**, Thumbnails shows selected times of three appearing exocytosis spots in the areas selected and represented in squares of 3 different colors green, red, and yellow in **(a). c**, The graphs represent the lifetime of the three same spots by measuring intensity. Note the behavioral differences of the red curve compared to the green and yellow ones. **d**, MATIRF image reconstruction^20^ was implemented in BioImageIT. MATIRF reconstruction parameters: depth: 300 nm, number of planes: 20, regularization: 0.001, numbers of iterations: 100. **e**, Representation of the first spot (Green square in (**a**)) reconstructed in 3D with three time points, pixels size in x, y, z (0.16 µm, 0.16 µm, 0.05 µm), using Napari for visualization. Scale bar: 5µm.

## Acknowledgements

This work was supported by the France-BioImaging Infrastructure (French National Research Agency - ANR-10-INBS-04-07, “Investments for the future”). We acknowledge the Cell and Tissue Imaging (PICT IBiSA, Institut Curie) platform, also member of the national infrastructure France-BioImaging (ANR-10-INBS-04-01) for maintaining the spinning-disk confocal microscope.

## Author contributions

S.P. conceived the idea in discussions with J.S., C.K., C.A.V-C. and L.L.; S.P. developed the algorithms and implemented the software; C.A.V-C. and L.L. performed all experiments and reconstructions; J.S., C.A.V-C. and L.L. designed and implemented the analysis workflows; L.M. implemented algorithm wrappers and the software packaging; J.S. and C.K. supported and supervised the project; and S.P., J.S., C.K. and C.A.V-C. wrote the manuscript with input from the co-authors.

## Competing interests

The authors declare no competing interests.

## Additional information

**Supplementary information** is available for this paper at

**Reprints and permissions information** is available at

**Correspondence and requests for materials** should be addressed to J.S. or C.K.

## Methods

### Cell culture and fluorescence labelling

Hela cells stably expressing a CD-Mannose-6 Phosphate Receptor eGFP tagged construct (eGFP-M6PR small Ca++ dependent receptor) kindly provided by Dr B. Hoflack, (TU-Dresden, Germany) and Hela cells stably expressing eGFP–Rab5 were grown in DMEM-Glutamax medium supplemented with 10% FCS, as previously described^23^. For LLSM (Lattice Light Sheet Microscopy^11^) imaging, they were seeded 6 to 7 hours before image acquisition at 150.103 per well of a 6 well plate, containing 3 to 4 coverslips with a 5mm diameter, treated as previously described^11^. Coverslips were directly transferred to the lattice light-sheet microscope (LLSM) sample holder and inserted into the imaging chamber thermoregulated at 37°C containing 6 ml of LLSM medium (DMEM, without phenol red, with 1% BSA, 0.01% penicillin–streptomycin, 1mM pyruvate, and 20mM HEPES) containing the Tubulin TrackerTM DR (Tub-Tracker Deep Red, ThermoFisher Scientific) probe (Ex. 652nm/Em. 669) at a 1/2000 dilution of a stock solution of 20µg in 60µl DMSO. Image acquisition started 15 minutes after. The hTERT-immortalized RPE1 cells (Human Retinal Pigment Epithelial Cell) were purchased from ATCC (CRL-4000). RPE1 cells within 30 passage number were used and grown in DMEM-F12 medium without phenol red (Life Technologies, Carlsbad, CA) supplemented with 10% (vol/vol) fetal bovine serum (FBS) at 37°C in humidified atmosphere with 5% CO2. RPE1 cells were transiently transfected with plasmid coding for TfnR-pHluorin^24^, using Lipofectamin 3000, following basically the manufacturer recommendations (Invitrogen, Waltham, MA). Briefly, we used 2 μg of DNAs, completed with 125 μL with OptiMEM (Invitrogen, Waltham, MA) and 4µl of Reagent 3000 and mixed with 3.75 μL Lipofectamin 3000 completed with 125 μL with OptiMEM. The mixed solution was then incubated for 15-20 min at room temperature and finally added to cells grown, in 2ml culture medium on glass bottom petri dish at 80% confluency, for a further overnight incubation at 37 °C. When specified, RPE1 cells stably expressing mCherry-Lifeact^25^ were imaged with spinning disk microscope. They were cultured as described above for wild type RPE1 and were plated 5 to 6 hours before acquisition at 200-250.103 per glass bottom 35 mm diameter Petri dishes (Ibidi GmBH, Gräfelfing, Germany).

### Lattice light-sheet microscopy

LLSM was done on a commercialized version of a previously described setup^11^ from 3i (Denver, USA). Cells were scanned incrementally through a 20 μm long light sheet in 600 nm steps using a fast piezoelectric flexure stage equivalent to ∼325 nm with respect to the detection objective and were imaged using a sCMOS camera (Orca-Flash 4.0; Hamamatsu, Bridgewater, NJ). Excitation was achieved with 488- or 642-nm diode lasers (MPB Communications, Canada) at 10 % acousto-optic tunable filter transmittance with 300 mW and 100 mW respectively (initial box power) through an excitation objective (Special Optics 28.6× 0.7 NA 3.74-mm water-dipping lens) and detected via a Nikon CFI Apo LWD 25× 1.1 NA water-dipping objective with a 2.5× tube lens with a final pixel size of 104 nm. Lattice light-sheet imaging was performed using an excitation pattern of outer NA equal to 0.55 and inner NA equal to 0.493. Composite volumetric datasets were obtained using ∼5-ms exposure/slice/channel at a time resolution of 0.833 s per total cell volume (about 40 slices). One hundred time points were acquired within 1 to 2 minutes. Acquired data were deskewed, a necessary step to realigned image frames, using pycudadecon, a python wrapper for cudaDecon, which is a CUDA/C++ implementation of an accelerated Richardson Lucy Deconvolution algorithm^12^ (copyright to T. Lambert, Harvard Medical School, Boston, USA; https://github.com/tlambert03/pycudadecon). Then deskewed images were deconvolved also using pycudadecon. Later, tracking was performed using Trackmate^13^. Napari^14^, a multi-dimensional image viewer for Python (https://github.com/napari/napari), was used for 3D rendering. Full workflow, including Pycudadecon, Trackmate^13^ and Napari^14^, was implemented in BioImageIT. Rab5 stably expressing Hela cell was scanned incrementally through a 20 μm long light sheet in 500 nm steps using a fast piezoelectric flexure stage equivalent to ∼270 nm with respect to the detection objective and were imaged using a sCMOS camera (Orca-Flash 4.0; Hamamatsu, Bridgewater, NJ). Excitation was achieved with a 488-diode laser (Coherent, Santa Clara, CA) at 15 % acousto-optic tunable filter transmittance with 300 mW. Volumetric datasets were obtained using ∼10-ms exposure/slice at a time resolution of 2s per total cell volume (about 50 slices). One single volume was used for the analysis. Deskew was performed using pycudadecon prior denoising. LLSM Deskewed 3D noisy data could also be directly treated using Noise to Void (N2V)^22^ as a training-free denoising methods embedded in BioImageIT and as such could be applied as another preprocessing step in a workflow when other methods cannot.

### Multiangle Azimuthal Averaging TIRFM

Acquisition and high-resolution three-dimensional reconstruction of Multiangle Stacks were carried out as previously published and using the same setup^20^, except that the custom C++ code based on the CImg library (https://github.com/dtschump/CImg) used for reconstruction, was integrated in BioImageIT. Our current setup is based on a Nikon Ti Eclipse equipped with the Ring-TIRF ILas2 module (GATACA systems, Massy, FR), with a PLAN APO TIRF 100X oil objective (1.49 NA). The acquisition of multiangle TIRF image stacks is performed from the iLas2 software interacting with the Metamorph 8.6 software (Molecular Devices, San Jose, CA). This system is interfaced with a hardware and software implementations allowing the full control of the illumination incident angle as a function of penetration depth for multiple wavelengths. TfR-Phluorine was imaged with excitation at a wavelength of 488nm. Nominal laser powers were adjusted at 10% for the 488nm laser (100 mW). The time-gated fluorescence detection was done an Evolve 512 (Teledyne Photometrics, Tucson, AZ) EMCCD camera. Readout time was down to 30 ms per frame.

### Fast confocal microscopy

It was performed with a spinning disk confocal based on a Ti-Eclipse inverted microscope (Nikon, Tokyo, JP) equipped with X1 confocal head (Yokogawa, Tokyo, JP), Prime95B, an sCMOS camera (Teledyne Photometrics, Tucson, AZ) and interfaced with a super resolution Live-SR module (GATACA systems, Massy, FR) based on optically demodulated structured illumination technique. Cells were scanned incrementally through ∼10 μm Z-stacks in 300 nm step at 30ms/frame, using a 100X, Plan Apo NA 1.4 oil objective (Nikon, Tokyo, JP). Excitation was achieved at 561-nm (50 mW), diode laser (GATACA systems, Massy, FR), at low illumination of the sample adjusting the acousto-optic tunable filter transmittance at 20 % in the Live-SR mode, allowing relatively long-term imaging without photobleaching and phototoxicity at high resolution. The acquisition and reconstruction of the Live-SR module is performed with the Metamorph 8.6 software (Molecular Devices, San Jose, CA). Reconstruction of resulting low signal to noise ratio (SNR) super resolution images was performed directly using the setup-software before denoising using the ND-SAFIR 4D software ^21^ implemented in BioImageIT.

## Data availability

Original datasets are available for download at https://github.com/bioimageit/manuscript.

## Code availability

All source code used in this publication is open-source and published under the BSD 4-Clause. The latest stable releases used in this publication and the current versions that include bugfixes and updates can be downloaded from GitHub (https://github.com/bioimageit). Details on how to use the software are described in website of the project https://bioimageit.github.io/.

## Supplementary Figure, Tables, Notes

Note:

Supplementary Videos 1–5 are available in https://bioimageit.github.io/#/videos

Supplementary Note 1 is available in https://bioimageit.github.io/bioimageit_core

Supplementary Note 2 is available in https://bioimageit.github.io/bioimageit_gui

## SUPPLEMENTARY FIGURE

**Supplementary Figure 1.**
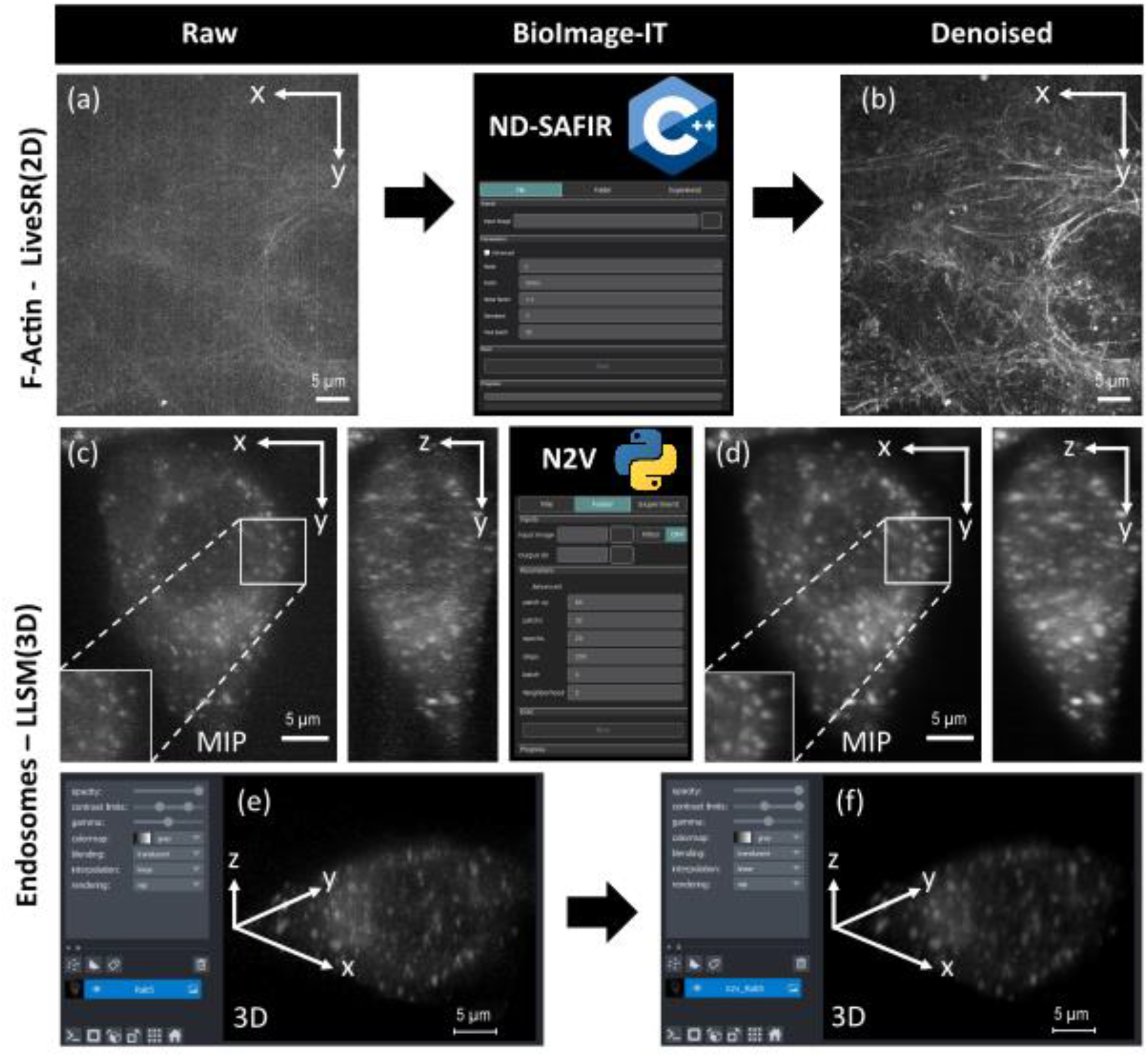
BioImageIT for advanced image optimization and Deep Learning. **a, b** Example of 2D-denoising using a conventional approach (ND-SAFIR^1^) of RPE1 cells stably expressing mCherry-LifeAct. (a) Raw Live-SR images. **b**. ND-SAFIR Denoised image. ND-SAFIR denoising was implemented in BioImageIT. ND-SAFIR parameters: Patch: 9×9×3, Noise factor: 1.3, Iterations: 5 and Noise type: Gaussian. **c-f**, Example of 3D denoising using a Deep Learning approach (Noise2Void^2^). 50 planes 3D volumes of live Hela cells stably expressing EGFP-Rab5 were acquired within 2 s per stack using lattice light-sheet microscopy^3^. MIP of representative raw images for endosomes (Rab5), before **(c)** and after **(d)** Noise2Void 3D denoising. Insets are zoomed area illustrating Noise2void improvement in SNR. 3D angular views before **(e)** and after **(f)** Noise2Void treatment. 3D-denoise parameters: patch xy=64, patch z=32, epochs=20, steps=200, batch=4 and neighborhood=5. Noise2void was applied to deskewed images as indicated in Figure 1. One single volume was used as input and target during the training.

## Description of Additional Supplementary Files

<u>Supplementary Video 1:</u>

Movie of installation of BioImageIT in a computer with Windows 10 operating system.

<u>Supplementary Video 2:</u>

Movie of installation of BioImageIT in a computer with Linux Ubuntu 21.04 operating system.

<u>Supplementary Video 3:</u>

Movie of installation of BioImageIT in a computer with Mac OS Catalina 10.15.11 operating system.

<u>Supplementary Video 4:</u>

Movie showing the workflow of ND-SAFIR^1^ applied to Genome-edited Hela cell (CD-M6PR-eGFP) 3D stacks acquired within 4.3 s per stack, exposure time: 20 ms, z-planes: 22, z-step: 0.4µm, 55 time points using spinning disk confocal microscopy.

<u>Supplementary Video 5</u>

Movie showing how to import and annotate data in BioImageIT.

Supplementary Note 1:

This file includes the developer manual. This developer manual was generated using Sphinx^4^. Code source can be found in https://github.com/bioimageit/bioimageit_core/tree/master/docs

Supplementary Note 2:

This file includes the user manual. This user manual was generated using Sphinx^4^. Code source can be found in https://github.com/bioimageit/bioimageit_gui/tree/master/docs

## Notes

https://bioimageit.github.io

### Competing Interest Statement

The authors have declared no competing interest.

